# Ultra fast tissue staining with chemical tags

**DOI:** 10.1101/005298

**Authors:** Johannes Kohl, Julian Ng, Sebastian Cachero, Michael-John Dolan, Ben Sutcliffe, Daniel Krueger, Shahar Frechter, Gregory S.X.E. Jefferis

**Affiliations:** Division of Neurobiology, MRC Laboratory of Molecular Biology, Cambridge, CB2 0QH, UK

## Abstract

Genetically encoded fluorescent proteins and immunostainings are widely used to detect cellular or subcellular structures in thick biological samples. However, each approach suffers from limitations, including low signal and limited spectral flexibility or slow speed, poor penetration and high background, respectively. Here we overcome these limitations by using transgenically expressed chemical tags for rapid, even and low-background labeling of thick biological tissues. We construct a platform of widely applicable transgenic *Drosophila* reporter lines, demonstrating that chemical labeling can accelerate staining of whole-mount fly brains by a factor of 100×. Together, this tag-based approach drastically improves the speed and specificity of labeling genetically marked cells in intact and/or thick biological samples.

## INTRODUCTION

The revolution in live imaging due to genetically encoded fluorescent proteins (FPs) is widely appreciated ^1, 2^, but FPs have also had a major impact on studies of fixed whole mount specimens or thick sections. Here immediate visualization, low background and spatially even signal are major advantages. However, FP signals are typically weaker, easily quenched by fixation and suffer from limited spectral flexibility. Therefore, antibody detection of marker proteins remains essential in many experimental situations. This tradeoff between uneven, slow staining and comparably high background levels due to antibody approaches, and the convenient but weaker, more labile and spectrally less diverse signals obtainable with FPs is central to many recent technical developments (e.g. array tomography^3^, CLARITY^4^, Sca*l*e^5^, SeeDB^6^ and CUBIC^7^), but remains fundamentally unresolved.

This tradeoff is also a major practical issue in neural circuit tracing in *Drosophila* ^8–13^. We therefore sought staining methods that combine the positive aspects of both FP and antibody-based staining, notably fast, even, strong signal vs. low-background labeling. We have developed an approach based on four commercially available, orthogonal labeling chemistries (SNAP-, CLIP-, Halo- and TMP-tag) characterized by the covalent binding of a large range of (fluorescent) substrates to engineered enzyme tags. In order to use these chemistries for effective tissue labeling, we have generated the first stable transgenic reporter animals bearing these tags. We validate their use and expression in *Drosophila*, presenting 12 widely applicable fly strains for labeling of cells and sub-cellular structures. Combining these tags with the first knock-in construct, we demonstrate that our approach can speed up the most widely-used antibody staining procedure in *Drosophila* neurobiology ^12, 14^ from 1 week to 1 hour, a factor of > 100, while giving more even staining and reduced background signals.

In conclusion, the chemical labeling reagents that we have developed and validated solve a basic but pervasive problem in tissue labeling and have immediate applications across model organisms and experimental disciplines.

## RESULTS

### Expression of chemical tags in the *Drosophila* brain

We sought to develop a labeling system that overcomes the limitations of antibody-based immunostainings, i.e. speed (poor penetration of thick tissue samples), specificity (background staining due to off-target binding) and complexity (number of user interactions, i.e. manual steps, in staining protocols).

Existing chemical tagging systems were compared and four were chosen that (1) do not require co-factors, (2) result in formation of a covalent bond and (3) for which a commercially available, spectrally diverse, range of fluorescent substrates is available. SNAP- and CLIP-tag ^15–18^ (NEB), Halo-tag ^19, 20^ (Promega) and TMP-tag ^21–23^ (Active Motif) fulfilled these requirements. All four rely on the rapid (10^2^ to 10^6^ M^−1^ s^−1^) formation of a covalent bond between an engineered, bio-orthogonal enzyme (tag) and a small reactive group which is fused to a reporter (substrate) ^24^. SNAP- and CLIP-tag are modified, 181 amino acid (aa) O^6^-alkylguanine-DNA alkyltransferases that bind benzylguanine (BG) or benzylcytosine (BC) derivatives, respectively. Halo-tag (295 aa) is an engineered bacterial haloalkane dehalogenase that binds chloroalkane groups and TMP-tag (159 aa) is an *E. coli* dihydrofolate reductase (eDHFR) which has been engineered to covalently bind trimethoprim (TMP) derivatives (see Fig. 1a).

**Figure 1:**
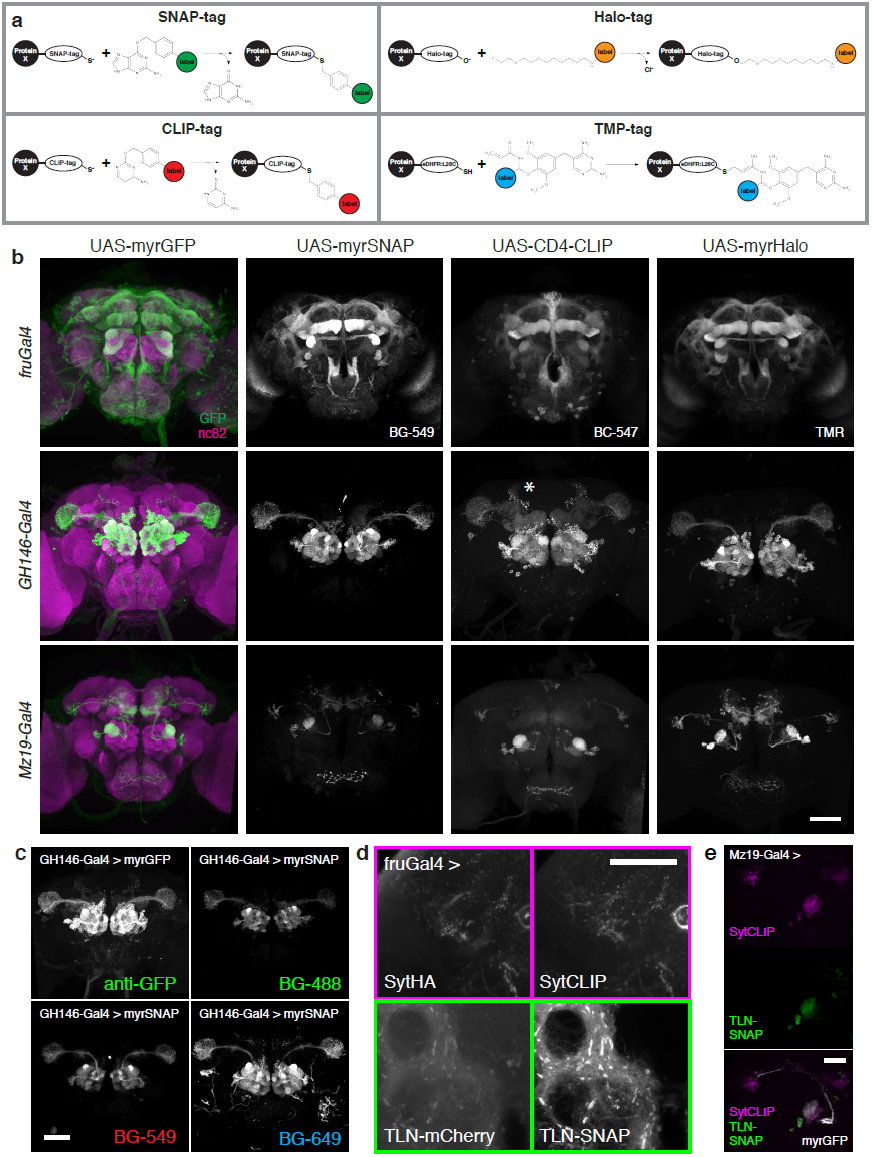
Expression of chemical tags in the fly brain. **(a)** Chemical labeling chemistries used in this study ^15, 16, 19, 23^; see text for description. **(b)** Expression patterns of membrane-targeted tags in the *Drosophila* central brain. The panel is arranged in a 3 row × 3 column grid. Rows represent Gal4 driver lines, columns represent reporter constructs. In the first column, nc82 neuropil counterstaining is shown (magenta). Fluorescent substrates used are indicated for each tag. Note that while *myrGFP*, *myrSNAP* and *myrHalo* are targeted attP insertions, *CD4-CLIP* is a P element insertion. Due to positional effects, the *CD4-CLIP* reporter shown here labels the bilateral APL neuron when crossed to *GH146-Gal4* (asterisk). **(c)** Brains of *GH146-Gal4 > myrSNAP*, *myrGFP* animals stained with GFP antibody (top, left), BG-488 substrate (top, right), BG-549 substrate (bottom, left) or BG-647 substrate (bottom, right). **(d)** Comparison between established synaptic markers and synaptically targeted chemical tags. Confocal slices from the brains of a *fru^Gal4^> SytHA, SytCLIP* (presynaptic markers, top) and a *fru*^Gal4^*> TLN-mCherry (DenMark), TLN-SNAP* (somatodendritic markers, bottom) animal are shown. **(e)** Simultaneous labeling of presynaptic sites and the somatodendritic compartment of *Mz19* DA1 projections neurons. Scale bars, 50 µm. See Table S2 for abbreviations of substrates.

We made fusion constructs of these tags with membrane- or cytosolic proteins and cloned them downstream of a Gal4 UAS sequence ^25^. The constructs were then injected into *Drosophila melanogaster*, generating a total of 12 transgenic lines (see Table S1).

We tested expression of these new fusion constructs using three different Gal4 driver lines: *fru*^Gal4^, which drives expression in about 2,000 *fruitless*-positive neurons ^26^, *GH146-Gal4*, which expresses in most olfactory projection neurons ^27^ and *Mz19-Gal4* which is expressed in three classes of projection neurons, DA1, VA1d and DC3 ^28^. As a reference, each driver line was crossed to flies expressing membrane-targeted GFP (myrGFP). When brains from the progeny of these driver-tag crosses were incubated with fluorescent SNAP-, CLIP-, Haloor TMP substrates (see Table S2), strong, specific labeling was observed (Fig. 1b and Fig. S1). We tested all tag-substrate combinations for potential cross-reactivity and found no signal from non-cognate tag-substrate pairs, with the exception of weak binding of CLIP substrates to SNAP-tag (Fig. S2). This is expected since it has previously been shown that there is a ∼100-fold preference of BC substrates for CLIP (an engineered version of SNAP) over SNAP, as opposed to a > 1,000-fold preference of BG groups for SNAP over CLIP) ^16^. Because of this low cross-reactivity and the availability of spectrally diverse substrates (Fig. 1c), this labeling approach potentially allows the simultaneous visualization of up to four orthogonal channels.

**Figure 2:**
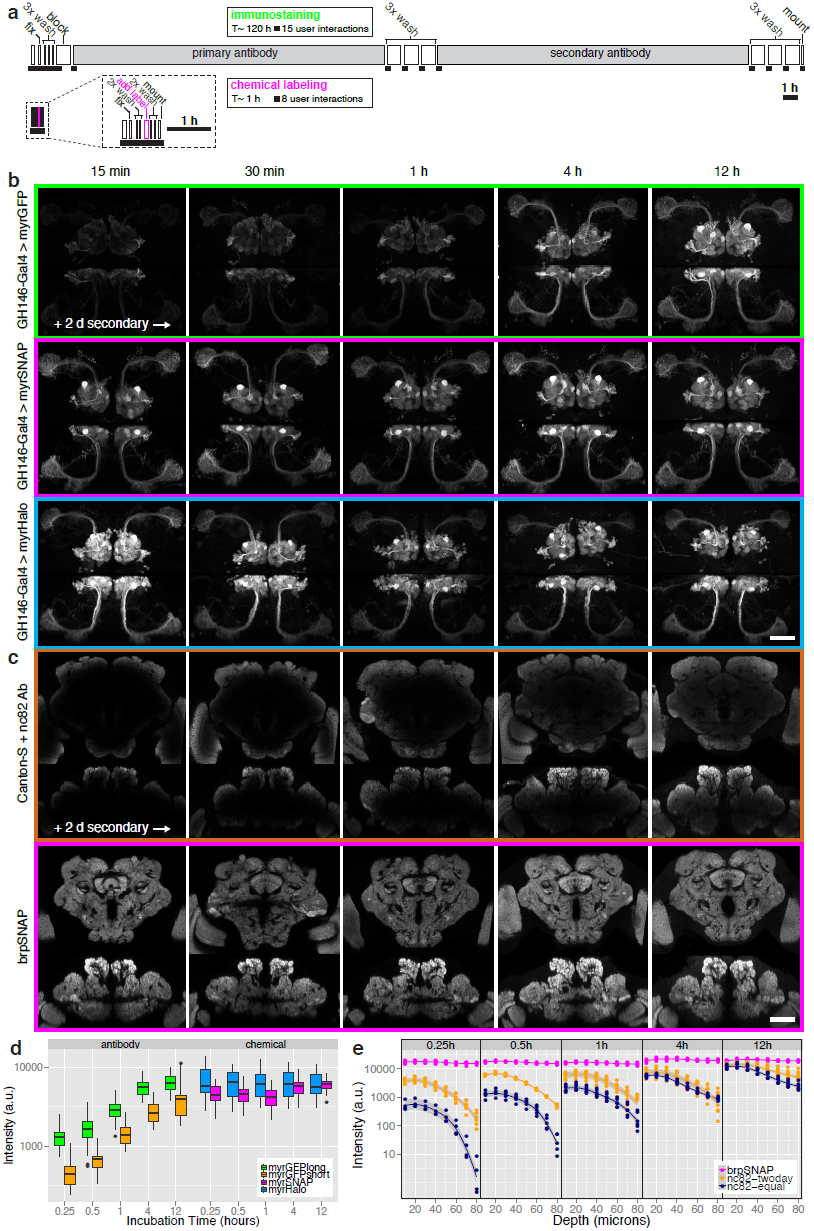
Ultrafast and homogeneous tag-based tissue staining. **(a)** Direct comparison of immunostaining and chemical labeling protocols. In these ‘ethograms’ the length of individual steps is proportional to the time required, and each rectangle represents a manual interaction. Chemical labeling is > 100× faster (∼1 h vs. > 100 h) and requires half as many (8 vs. 15) manual handling steps of the sample. **(b)** Staining time course of *GH146* projection neurons using immunostaining against membrane-targeted GFP (top row), chemical labeling using SNAP-tag (middle row) or Halo-tag (bottom row). *z*- and *y-* maximum intensity projections from 3D confocal stacks are shown, incubation times are indicated for primary antibody or chemical substrates (GFP antibody, BG-549, Halo-TMR, nc82-antibody). Note that incubation with secondary antibodies was for 2 days. **(c)** Staining time course of the nc82/Bruchpilot neuropil marker using immunostaining against Brp protein (top row) or chemical labeling using SNAP-tag (bottom row). Single coronal and horizontal confocal slices through the center of the brain are shown. Note that incubation with secondary antibodies was for 2 days. **(d)** Quantification of signal intensity over time in *GH146-Gal4* brains labeled with antibody vs. chemically. Based on brains displayed in (b) and Fig. S5a. Fluorescence was quantified using a *GH146* mask (see Fig. S7b and Methods, n = 5–8 brains per condition). **(e)** Quantification of labeling intensity (see Methods) at different depths from the brain surface in brains labeled with nc82 antibody vs. chemically (using *brp-SNAP*). Secondary antibodies for nc82 immunostainings were either incubated for 2 d (’nc82-twoday’) or for the same duration as primary antibodies (’nc82-equal’). Five different time points are shown. Scale bars, 50 µm.

Having demonstrated efficient plasma membrane labeling, we then constructed synaptically localized tags, comparing them with established synaptic markers in *Drosophila*. Simultaneous immunostaining against the synaptotagmin-HA presynaptic marker (SytHA ^8, 29, 30^) and chemical labeling with a SytCLIP construct revealed that chemical labeling consistently achieved higher signal-to-noise ratios with identical marker localization (Fig. 1d, top). This was even more pronounced when the somatodendritic marker TLN-mCherry (DenMark ^31^) was compared to a TLN-SNAP construct (Fig. 1d, bottom). We used these synaptic markers to simultaneously label pre- and postsynaptic compartments of DA1 projection neurons with high signal-to-noise ratio (Fig. 1e). This highlights that by being bio-orthogonal, the chemical labeling approach can overcome one of the most common shortcomings of immunohistochemistry: high background staining due to poor epitope-specificity and/or cross-reactivity.

So far we have demonstrated (a) that chemical tags can be efficiently and specifically expressed in the *Drosophila* brain, (b) that these tags can be simultaneously used thanks to their orthogonality and the spectral range of their fluorescent substrates and (c) that the signal-to-noise ratio of this chemical labeling approach is superior to that commonly achieved with antibody staining.

### Ultrafast labeling of thick tissue samples

Low fluorescence after fixation is a widely appreciated limitation of genetically encoded FPs ^7, 32, 33^. Indeed, we observed that most fluorescence was quenched in *GH146-Gal4 > myrGFP* brains after standard fixation with PFA (Fig. S5f). Therefore, immunostainings are usually required to obtain sufficient signal from fixed samples. Antibody diffusion into tissue is another rate-limiting step for immunostainings of thick specimens. For example, homogeneous immunostaining of *Drosophila* brains (approximately 500 × 250 × 200 µm in *x*, *y* and for the nc82/Bruchpilot synaptic protein ^34^ (the standard counterstain for brain structure in *Drosophila* neuroanatomical studies, see below) takes seven days (Fig. 2a) and requires > 15 user interactions. This comprises a blocking step and prolonged incubations with primary and secondary antibodies ^9, 12^. These long staining times are due to the relatively large size of antibodies (∼150 kDa), which limits their diffusion into thick tissue samples. We reasoned that the much smaller size (∼1 kDa) of fluorescent chemical labeling substrates would considerably speed up the staining process.

We directly compared the labeling speed of antibodies and chemical substrates in fly brains expressing either membrane-targeted GFP or membrane-targeted SNAP-tag in olfactory projection neurons (PNs). Brains were dissected, fixed and briefly permeabilized, then incubated with GFP antibody or fluorescent SNAP substrate for 15 min, 30 min, 1 h, 4 h or 12 h (see Methods). After a washing step, fluorophore-conjugated antibody was added to brains expressing GFP for 2 d. Brains expressing SNAP-tag were briefly washed and directly mounted. In a separate time series experiment, the incubation times in primary and secondary antibodies were identical (Fig. S5a). Whereas homogeneous staining of PNs with anti-GFP antibody required > 4 h primary antibody followed by 2 d of secondary antibody (Fig. 2b, top), strong and uniform chemical labeling was visible after only 15 min (Fig. 2b, middle). Such rapid labeling was also observed when using membrane-targeted Halotag (Fig. 2b, bottom). Quantification of staining intensity (see Methods) revealed that while antibody staining increased over the course of 12 h, chemical staining levels reached near-maximum levels after 15 min (Fig. 2d). These results show that the small substrate size indeed leads to drastically reduced staining times with chemical labeling. Because no blocking step and far fewer washing steps were required for chemical labeling, the protocol comprised half as many user interactions (Fig. 2a).

We next assessed labeling uniformity and speed for the nc82/Bruchpilot antigen ^34^; we compared nc82 antibody labeling of wild-type (Canton-S) brains with labeling of a SNAP-tag knock-in construct inserted into the *brp* locus (*brp-SNAP*, see Fig. S4). Strikingly, while nc82 immunostaining for 12 h (primary) followed by 2 days (secondary) still resulted in substantial signal drop-off in the center of the brain (Fig. 2c, top), chemical labeling produced uniform staining within 15 min (Fig. 2c, bottom). Indeed we found that staining times of only 1 min were sufficient (Fig. S5d). Longer incubations did not appreciably increase neuropil labeling but instead resulted in higher background (visible in Fig. 2c, bottom right panel); this non-specific labeling could be removed by extended washes (see Fig. S6). This indicates that most tag molecules are occupied by substrate after minutes of incubation. We directly tested this by sequential labeling of the same sample with two different substrates (Fig. S6).

Using immunohistochemistry, we found that labeling the center of the brain (∼100 µm from surface) required at least 12 h of primary followed by 2 d of secondary antibody incubation (Fig. 2e). In contrast, fluorescent substrates uniformly labeled the sample within less than 15 min (Fig. 2e). Therefore, the chemical staining approach enables rapid and homogeneous staining of thick tissue samples.

Finally, we tested the effect of fixation on the labeling reaction, incubating samples for 20, 40 or 60 min in 4% paraformaldehyde before adding substrates. We found that while SNAP- and CLIP-tag were largely insensitive to prolonged fixation, Halo-tag labeling decreased and TMP-tag labeling increased with longer fixation (Fig. S3). We did not observe fixation-dependent changes in background labeling (Fig. S3 and data not shown).

**Figure 3:**
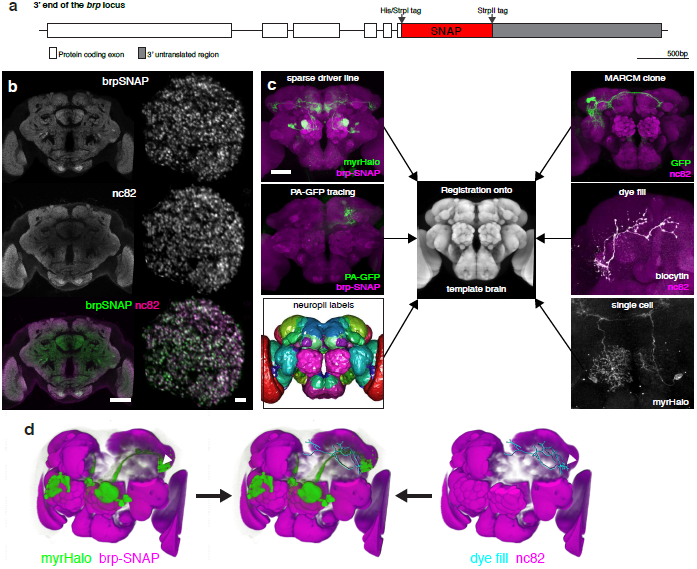
Key applications for chemical labeling in *Drosophila* neurobiology. **(a)** Map of the *brp-SNAP* knock-in. **(b)** nc82 and *brp-SNAP* signals co-localize in a *brp-SNAP/+* brain simultaneously labeled with BG-549 and nc82 antibody. Single coronal slices through the middle of the brain (left) and through a deconvolved image stack of the DA1 glomerulus (right) are shown. Note the more even *brp-SNAP* staining in the center of the brain. Scale bars, 50 µm (top), 2 µm (bottom). **(c)** Chemically labeled neurons can be successfully registered onto a template using the *brp-SNAP* neuropil counterstaining. This allows direct comparison with neurons from other sources, such as stochastic labeling (MARCM), PA-GFP tracing experiments, whole-cell recordings or sparse driver lines (the *Mz19-Gal4* expression pattern and a single neuron from *GMR14C11-Gal4* are shown). Scale bar, 50 µm. **(d)** Overlay of DA1 projection neurons (green) from a chemically labeled *brp-SNAP, Mz19-Gal4 > myrHalo* brain (left) overlaid with a dye-filled third-order olfactory neuron (cyan) from an nc82-stained brain (right) after registration. Notice the overlap between axon terminals and dendritic arbor (white arrows).

These results show that chemical labeling of thick tissue samples is at least two orders of magnitude faster than immunostainings and requires half as many user interactions, while at the same time resulting in better tissue labeling (Fig. 2a and Table 1).

**Table 1:** Comparison of immunostainings with chemical labeling

|  | **immunostaining** | **chemical labeling** |
| --- | --- | --- |
| **duration** | hours-days | minutes-hours |
| **user interactions** | > 15<sup>(a)</sup> | 8 |
| **substrate size** | ~150 kDa | ~1 kDa |
| **staining steps** | 2<sup>(b)</sup> | 1 |
| **blocking** | Yes | No |
| **background** | often high | low |
| **epitope stability** | variable | high |
| **cell permeable** | No | Yes/No |
| **No. of channels** | many | 4 (currently) |
| **transgenics** | No | Yes |
| **stoichiometry<sup>(c)</sup>** | > 1:1 | 1:1 |

Comparing antibody-based and tag-based staining approaches. Staining durations indicated are typical for thick tissue samples (hundreds of μm to several mm). Notes: (a) Using standard immunostaining protocol for Drosophila brains (see text for references). (b) Fluorophore-coupled primary antibodies are sometimes used, reducing the number of staining steps to 1. (c) i.e. label-to-tag ratio.

### Applications in *Drosophila* Neurobiology

The *Drosophila* reagents that we have described here have immediate and widespread utility in cell, developmental and neurobiology. In order to demonstrate this more clearly we optimized *Drosophila* transgenics for applications that are at the heart of large scale studies of neural circuits in the fly brain. The nc82 marker is the most widely used counterstain for the *Drosophila* brain (currently 753 publications listed in Google Scholar http://scholar.google.co.uk/scholar?q=%22nc82%22+AND+%28monoclonal+OR+antibody%29&btnG=&hl=en&as_sdt=0%2C5). It has been used to construct standard brains for 3D atlases by image registration ^8, 9, 12, 35^ and is the basis of a 3D model encapsulating the recently standardized nomenclature for *Drosophila* brain regions ^36, 37^ (see also http://www.virtualflybrain.org). Homogeneous staining of neuropil structures throughout the brain is absolutely critical for successful registration ^9, 38^.

We first verified that the new *brp-SNAP* knock-in (Fig. 3a) closely recapitulates the neuropil staining obtained with nc82 antibody (Fig. 3b). We then confirmed that chemically labeled brains can easily be registered onto an existing nc82-immunostained template brain (IS2 ^9^) with 100% success (n = 84 specimens). Therefore, using the chemical *brp-SNAP* counterstain, rapidly labeled brains from different experimental sources can be comfortably registered into the same reference space, enabling direct comparison of labeled structures. In order to demonstrate this we generated a fly line bearing both a *UAS-myrHalo* reporter gene and *brp-SNAP,* crossed it to the sparse *Mz19-Gal4* line and registered the brains of progeny after 30 min of labeling (Fig. 3c). We were subsequently able to resolve fine details of individual neurites and presynaptic boutons in the labeled *Mz19-Gal4* expression pattern (Fig. 3c). Even when crossed to comparatively weak driver lines our chemical labeling reporter lines gave strong enough signal to visualize single neurons (Fig. 3c). Another powerful technique for sparse labeling of neuronal subsets within a wider expression pattern is the use of photoactivatable GFP (PA-GFP); here spatially targeted laser stimulation can identify neurons with processes in a specific region of the brain ^39, 40^. However, although these methods effectively reveal neuronal morphology, they cannot easily be used with image registration approaches since the PA-GFP signal does not survive standard neuropil immunostaining protocols. We demonstrate that *brp-SNAP* labeling can be used to substitute for nc82 labeling in this context as well, allowing co-registration of PA-GFP specimens (Fig. 3c).

All of these newly generated data can be integrated with existing resources defined by registration against nc82 template brains, such as stochastically labeled single neurons ^38^, lineage clones ^9, 41, 42^, large-scale expression screens ^12^, single recorded neurons ^13^ or the standard neuropil regions defined by the Insect Brain Name working group ^37^ (Fig. 3c). We demonstrated this important point by co-registration of chemically labeled *Mz19-Gal4* DA1 projection neurons with a dye-filled third-order olfactory neuron ^13^ (Fig. 3d).

## DISCUSSION

This work aims to address a fundamental need for visualizing genetically marked cellular structures in thick or intact tissue samples with high signal-to-noise ratio. It combines advantages of immunostainings (spectral flexibility, high signal after fixation) and genetically encoded fluorophores (low background) while drastically reducing staining time. The approach presented here is practical: it utilizes extensively tested and commercially available reagents in a novel context. Thus, little or no optimization will be required by researchers wishing to introduce this method. Because the building blocks for coupling reporter molecules to reactive groups are also available, customized substrates can be made by individual groups. Use of suitable probes should allow to combine chemical labeling with electron or super-resolution microscopy in the future. One advantage of chemical labeling in this context is its inherent linearity (i.e. one substrate molecule binds to one tag molecule), in contrast to often highly non-linear immunostainings. Chemical labeling of appropriate cellular targets is therefore especially suited for quantitative studies.

While our approach requires the use of transgenic animals and therefore can not replace the use of antibodies against specific cellular proteins, recent advances in molecular biology ^43–46^ have greatly facilitated the generation of even complex DNA constructs. Furthermore, the CRISPR method now enables the very rapid generation of transgenic animals at defined loci, either for introducing reporter genes or tagging endogenous proteins ^47–53^.

Our experiments show that SNAP-, Halo- and CLIP-tags are suitable for single-cell labeling. In contrast, using TMP-tag for this purpose will require further optimization. Other, orthogonal, labeling chemistries such as ACP-tag ^54^ could be explored in the future to allow simultaneous visualization of > 4 channels. The quantitative nature of chemical labeling also makes it particularly suitable for multiplexing strategies such as Brainbow ^55^. Two Brainbow approaches are currently available in flies ^32, 56^. In our hands antibody staining against dBrainbow ^56^ proteins is too variable across the brain to enable segmentation of neurons by color; conversely, Flybow ^32^ fluorescent proteins have insufficient signal-to-noise ratio for mapping individual neurons in the central brain. The data in this paper suggest that chemical labeling would overcome these difficulties. We are currently working on such approaches for rapid, multicolor labeling of neuronal tissue in flies and mice.

We have generated transgenic fly lines suitable for a wide range of applications in cell and developmental biology (see Table S1). In the field of neurobiology, using membrane-targeted reporter lines in combination with *brp-SNAP* will considerably speed up anatomical studies where high resolution confocal imaging requires fixed and stained specimens. Furthermore, although intensity-based image registration has been used to great effect in large-scale anatomical studies ^8, 9, 11, 12^, it has not yet been widely adopted – one major factor being the difficulty of obtaining successful neuropil staining using nc82 antibody.

We have noticed that the limiting factor for spatially even immunostaining of thick samples is diffusion of secondary antibody into the tissue: while antibody-penetration into fly brains is incomplete after 12 h incubation with each primary and secondary antibody (Fig. S5b), prolonging secondary incubation to two days results in more homogeneous labeling (Fig. 2c, top). Indeed, others have recommended extending secondary antibody incubations to four days ^12^. Arguably, incubation times are less important in a high-throughput context. However, our approach requires half as many manual handling steps, resulting in proportional labor and cost savings. In any case, reducing a standard staining time from one week to one hour results in a shortening of the experimental cycle that has major benefits: conditions can be rapidly optimized and experimental decisions can be made faster. Furthermore, we find that inexperienced users can obtain even and robust labeling with this method whereas e.g. nc82 immunostaining is subject to batch-to-batch variation, even in experienced hands.

Rapid staining may be of even greater significance in larger specimens. For example several recent methods can render tissue sufficiently transparent to enable whole-mount optical imaging of mouse brains ^4–7^. However, high-resolution, multichannel imaging of fluorescent proteins has major limitations in sensitivity. The recently developed CLARITY technique additionally promises to enable antibody staining of intact brain tissue ^4^. Nonetheless, the reported staining time of four weeks for an adult mouse brain (∼5 mm across, i.e. 2.5 mm from each surface) will likely be prohibitive for many experiments and there are still signs of significant spatial inhomogeneity. Extrapolating from our results in ∼200 µm fly brains, chemical labeling could reduce this step to < 3 hours, a more than 200-fold improvement, while enabling more spatially even labeling. We are presently investigating the use of chemical labeling in mice.

## ONLINE METHODS

### *Drosophila* stocks

All flies used in this study were 2–4 d old males with the exception of PA-GFP tracing experiments (see below). *fruitless^Gal4^* (*fru^Gal4^*) is a targeted insertion of the yeast transcription factor Gal4 into the P1 promoter of the *fruitless* gene ^26^. *GH146-Gal4* ^57^ and *Mz19-Gal4* ^28^ are P element enhancer trap insertions. The generation of constructs for transgenic fly stocks is described below. *UAS-DenMark* ^31^, *Syt-EGFP* ^29^*, syb-EGFP* ^29^ and *myr-GFP* ^58^ flies were obtained from Bloomington Stock Center.

### *Drosophila* constructs

- pTW<Syt-CLIPm>, pTW<Syt-SNAPm>, pTW<CLIPf-Syb> and pTW<SNAPf-Syb> constructs were made using the Multisite Gateway technology platform (Invitrogen). CLIPf and SNAPf are engineered versions of the original versions CLIP26m (CLIPm) and SNAP26m (SNAPm), respectively, that display faster labeling kinetics ^17, 18^. pTW is a pUASt vector containing a Gateway cassette. All CLIPm, CLIPf, SNAPm and SNAPf coding sequences were amplified from pCLIPm, pCLIPf, pSNAPm or pSNAPf plamids (NEB), respectively. *Drosophila synaptotagmin 1 (syt1,* GenBank M55048) was amplified from *P{UAS-syt.eGFP}1* flies (Bloomington, ^29^) using *syt* forward and *EGFP* reverse primers. *Drosophila n-synaptobrevin* (*n-syb*, GenBank S66686) was amplified from *P{UAS-n-syb.eGFP}2* flies using *syb* forward and EGFP reverse primers. All synaptotagmin constructs described in this study are C-terminal, i.e. cytoplasmic fusions. All synaptobrevin constructs described here are N-terminal fusions, i.e. cytoplasmic.
- UAS-TLN-CLIPm and UAS-TLN-SNAPm constructs were generated as follows: First, fusion PCR was performed on a pUAST-TLN-cherry plasmid to remove the mCherry coding sequence and flanking linker sequences (pUAST-TLNΔcherry). A 1.5 kb fragment from the XhoI restriction site to the sequence corresponding to amino acids TVRVA of mouse Telencephalin/ICAM-5 and a 1.0 kb fragment ranging from amino acids GPWLW of Telencephalin/ICAM-5 to the MfeI restriction site were amplified. Using primers that introduced flanking XhoI and BglII sites, the full 2.7 kb fragment was cloned into pUAST-TLNΔcherry (pUAST-TLN). CLIPm and SNAPm sequences were amplified using BglII-containing primers and inserted into pUAST-TLN, thus yielding pUAST-TLN-CLIP and pUAST-TLN-SNAP.
- pTW<PAT3SP-CD4-CLIPf> and pTW<PAT3SP-CD4-SNAPf> constructs were made using the Multisite Gateway technology platform. PAT3 signal peptide (SP) and CD4 sequences were amplified from *UAS-CD4::spGFP1-10* flies ^59^.
- The UAS-myr-SNAPf construct was generated as follows: the GFP coding sequence was removed from a pJFRC-MUH-myr-GFP construct ^58^ using BamHI/KpnI restriction sites and the SNAPf coding sequence was amplified from pSNAPf (NEB) and inserted.
- The UAS-myr-TMP construct was generated as follows: the GFP coding sequence was excised from a pJFRC81-L21 construct ^60^ using BamHI/KpnI restriction sites and replaced with the eDHFR coding sequence (codon-optimized for *Drosophila*) and an N-terminal myristoylation signal.
- The UAS-myr-Halo construct was generated as follows: the TMP coding sequence was excised from UAS-myr-TMP (see above) using BamHI/KpnI restriction sites and replaced with a Halo-tag sequence amplified from the pHT2 Halo-tag plasmid (Promega).

### Generation of *brp-SNAP*

We targeted the *bruchpilot* (*brp*) gene taking advantage of a MiMIC ^61^ insertion (MI02987) 6.8 kb upstream of the stop codon present in the last coding exon of *brp* (Fig. S4). Because this exon is common to all nine Brp isoforms annotated in Flybase, we anticipated that inserting the SNAP ORF immediately 5' to the stop codon would label all Brp isoforms. The *brp-SNAP* targeting construct was generated by Gibson assembly ^44^ of the following fragments into the backbone of a pDONR221 plasmid.

- three fragments encompassing the DNA sequence from the MiMIC insertion site to the end of the last coding exon
- SNAP coding sequence (added 5' to the stop codon of *brp*)
- a 2.2 kb fragment of the *brp* 3' UTR and a cleavage site for the homing endonuclease I-CreI
- a fragment containing the 3xP3 promoter driving expression of RFP (used for screening of successful reduction step (see below).

The entire assembly was flanked by PhiC31 sites on both sides. Transgenic flies with the *brp-SNAP* construct replacing the original MiMIC insertion were made (BestGene Inc.) and genotyped to verify the landing site and orientation of the integration. Two positive lines were identified and assayed by anti-SNAP staining. Since the integration into the *brp* locus generates a large duplication and an incomplete 3' UTR (Fig. S4), one positive line was crossed to a *hs-I-CreI* line. The progeny were heat shocked for 30 min and subsequently crossed to balancers. Progeny from this cross were then screened for loss of the RFP eye marker, indicating successful repair of the I-CreI induced double strand break, resulting in reduction of the duplication and restoration of the 3' UTR. One such fly was identified and used to generate a stock wich was then verified by PCR (Fig. S4d) and anti-SNAP staining.

### Chemical stainings and immunohistochemistry

#### Labeling reagents

SNAP- and CLIP-Surface (i.e. cell-impermeable) labeling substrates are abbreviated as BG-(benzylguanine) or BC-(benzylcytosine) in this study (see Table S2 for all abbreviations). Substrates were either acquired as stock solutions (e.g. Halo-TMR) or acquired in powder form and dissolved in fresh, water-free dimethyl sulfoxide (DMSO, Sigma) at a concentration of 1 mM (SNAP and CLIP substrates). Aliquots (5 µl) were stored at –20 °C in presence of desiccant. We observed that using old DMSO or storing dissolved substrates in wet and/or warm conditions can lead to hydrolysis, drastically reducing labeling efficiency (see ‘Protocols for chemical labeling’ below).

#### Fly brains

Immunohistochemistry was carried out as follows: brains were dissected in 0.1 M phosphate buffer (PB), then fixed in 4% paraformaldehyde (PFA, Electron Microscopy Services, Hatfield, PA, USA) in PB at room temperature for 20 min (see ref [^8^]). We observed that prolonged fixation (1 h at room temperature) improved signal intensity for some far-red SNAP substrates (e.g. BG-TF5), while decreasing labeling of Halo-tag. For antibody stainings and non cell-permeable substrates, brains were then permeabilized by washing in PBT (phosphate-buffered saline + 0.3% Triton, 2 × 5 min). Cell-permeable substrates were added without permeabilizing the brains. For immunostainings, blocking with 5% goat serum was performed overnight at 4 °C. No blocking was performed for chemical stainings. All following washing steps were performed in PBT (immunostainings and chemical stainings with non cell-permeable substrates) or PBS (cell-permeable substrates) for 2 × 10 min. Fluorescent substrates (see Table S2) were added at a concentration of 0.1 − 1 µM for 1 min − 12 h (on a rotating wheel, at room temperature for incubations < 4 h, at 4 °C for incubations > 4 h). An increase in background signal was typically observed for incubations of cell-permeable substrates for > 2 h. For immunostainings, prolonged incubation of 2 days each with primary and secondary antibodies (rotating at 4 °C) was required for homogeneous staining. Primary antibodies were: mouse anti-nc82 ^62^ (DSHB, University of Iowa) 1:20–1:40, chicken anti-GFP (Abcam, ab13970) 1:1000, rat anti-HA (Roche, 11 867 423 1:200. Secondary antibodies (all from Life Technologies) were: Alexa-568 anti-mouse (A-11004) 1:1200, Alexa-633 anti-mouse (A-21052) 1:1200, Alexa-488 anti-chicken (A-11039) 1:1200, Alexa-568 anti-rabbit (A-11011) 1:1200. Specimens were whole mounted in Vectashield (Vector Laboratories) on charged slides to avoid movement.

### Image acquisition

Confocal stacks of fly brains were acquired using a Zeiss 710 confocal microscope. Brains were imaged at 768 × 768 pixel resolution every 1 µm (voxel size 0.46 × 0.46 × 1 µm) using an EC Plan-Neofluar 40×/1.30 Oil objective and 0.6 zoom factor. Images of dye-filled neurons were acquired with 2× (frame) averaging. Detail images were taken with a Plan-Apochromat 63×/1.4 Oil objective at 2–3× zoom and contained about 30 slices (768 × 768 pixels) with a voxel size of 0.06 × 0.06 × 0.15 µm. All images were taken using 16-bit color depths. Confocal images were registered to the IS2 template brain as previously described ^9^ using the Computational Morphometry Toolkit (CMTK) available at http://www.nitrc.org/projects/cmtk.

### PA-GFP tracing and brp-SNAP labeling

Brains from 1–2 day old flies were dissected in ice-cold artificial hemolymph (108 mM NaCl, 5 mM KCl, 2 mM CaCl_2_, 8.2 mM MgCl_2_, 4 mM NaHCO_3_, 1 mM NaH_2_PO_4_, 5 mM trehalose, 10 mM sucrose, 5 mM HEPES, pH 7.5, 265 mOsm) and allowed to adhere to a Poly-d-lysine treated coverslip within a cell culture dish. Photoactivation was performed with a two-photon microscope as described ^63^. Briefly, several Kenyon cells were selected for photoactivation using baseline fluorescence at 925 nm. The somata of these cells were continuously illuminated with 710 nm light for 2 min, followed by a 30 min resting period to allow for diffusion of photoactivated GFP. The brain was then fixed in 4% PFA in PB for 20 min at room temperature and washed in PBT (3×). Subsequently, 200 nM of BG-549 substrate in PBT were added for 20 min. Finally, brains were briefly washed with PBT, equilibrated in Vectashield, mounted and imaged.

### Image analysis and quantification of labeling

For quantification of substrate penetration into fly brains, samples from *GH146-Gal4 > myrSNAP* or *GH146-Gal4 > myrHalo* or *brp-SNAP* animals were incubated with 1 µM BG-549 or 1 µM Halo-TMR for 15 min, 30 min, 1 h, 4 h or 12 h. Samples from *GH146-Gal4 > myrGFP* or Canton-S animals were incubated with anti-GFP (1:1,000) or anti-nc82 (1:30) antibodies for 15 min, 30 min, 1 h, 4 h or 12 h, followed by washing and incubation with fluorescent secondary antibodies (anti-chicken Alexa 568 and anti-mouse Alexa 568, respectively) for either 2 d or for the same time as primary antibodies (i.e. 15 min, 30 min, 1 h, 4 h or 12 h).

#### Quantification of GH146 labeling

All 3D confocal stacks were registered onto a *GH146-Gal4 > myrGFP* template. GH146-positive areas were segmented from each registered brain using a binary mask (surfaced rendered in Fig. S7b) constructed as follows: five *GH146-Gal4 > myrGFP* brains (stained with anti-GFP antibody for 12 h) were averaged in Fiji. The resulting image stack was filtered (median + Gaussian) and thresholded.

#### Quantification of brp-SNAP/nc82 labeling

All images were registered against an nc82 template brain (IS2 ^9^). Two binary masks were made in order to separate neuropil and cortex (see Fig. S7a): five *brp-SNAP* brains labeled with BG-549 were averaged, filtered (median + Gaussian) and thresholded. In order to assess cortical background labeling, signal was quantified in the region resulting from subtraction of the neuropil mask from the whole brain mask (see Fig. S7a).

### Protocol for chemical labeling of *Drosophila* brains

- dissect brains in ice-cold 0.1 M PB
- fixation in 4% PFA (in 0.1 M PB) at room temperature for 20 min in glass-well plate on nutator

– SNAP-tag / CLIP-tag: in general no change in labeling intensity after prolonged fixation of up to 60 min however, we observed improved labeling intensity for BG-649 substrate after 60 min fixation.
– Halo-tag: labeling intensity significantly *decreased* with longer fixation (40–60 min)
– TMP-tag: labeling intensity significantly *increased* with longer fixation (40–60 min)
- (1) non cell-permeable substrates:

– transfer brains to 1.5 ml tube, permeabilize briefly (5–10 min) by incubation in 0.5 ml of PBT (0.3% Triton X-100) on rotating wheel
– add substrate for a final concentration of 1 µM in PBT (see notes on substrates below) higher concentrations can result in higher background
– incubate brains on rotating wheel for 15 min agitation of the sample(s) is important to obtain rapid, homogeneous staining! long incubation times (> 6 h) can result in increased background staining!
– wash brains with PBT (2 × 10 min)
- (2) cell-permeable substrates:

– transfer brains to 1.5 ml tube, wash briefly (5–10 min) in PBS
– add substrate for a final concentration of 1–5 µM in PBS higher concentrations usually result in considerable background; using cell-permeable substrates in presence of Triton typically results in very weak labeling (when using cell-permeable substrates on permeabilized samples, wash sample with PBS before adding substrate
– incubate brains on rotating wheel for 15 min agitation of the sample(s) is important to obtain rapid, homogeneous staining! long incubation times (> 6 h) can result in slightly increased background staining!
– wash brains with PBS, or – to reduce background – with PBT
- remove PBT as completely as possible and add ∼200 µl of Vectashield (or other) mounting medium we observed that subsequently transferring brains into a fresh ∼200 µl aliquot of Vectashield results in more homogeneous signal along *z*
- mount brains on charged slides and image

### Notes on substrates for chemical labeling

- store substrates at –20 °C in presence of dessicant this avoids hydrolysis of substrates
- for substrates that are purchased in dry powder form: dissolve in fresh, water-free DMSO using fresh DMSO from a sealed container is crucial as DMSO is hygroscopic; we observed reduced labeling efficiency when using substrates that were dissolved in old DMSO, presumably due to substrate hydrolysis
- avoid freeze-thawing, aliquot samples (e.g. 2 µl aliquots in 0.2 ml PCR tubes) we observed a decrease in labeling efficiency for some substrates that underwent several freeze-thaw cycles (e.g. BG-649)
- BG-488, BG-549, BG-649, BC-547, BC-488, Halo-TMR and TMP-Fluorescein work well using our standard staining protocol
- when using chemical labeling in combination with immunostainings, substrates should be added right after tissue permeabilization (not after the o/n blocking step) to obtain optimal results
- *Cost of chemical labeling:* the cost of 50 nmol of SNAP- or CLIP-substrate (sufficient for ∼1,000 labeling reactions at 1 µM concentration in a volume of 0.5 ml each) is comparable to that of an 0.5 ml aliquot of fluorophore-coupled secondary antibody (sufficient for ∼1,000 immunostainings in a volume of 0.5 ml each). However, we have successfully tested some substrates (e.g. BG-549, Halo-TMR) at 100 nM and lower concentrations (indeed, there might be situations where lower substrate concentrations – combined with longer incubation times – might be desirable). Also, no primary antibodies and blocking reagents are required for chemical labeling. Therefore, the cost of chemical labeling is comparable to, or even lower than, the cost of immunostainings.

## ACKNOWLEDGMENTS

We thank K. Johnsson, V. Cornish and I. Correa for providing labeling reagents, A. Carter and B. Hassan for DNA constructs, the Developmental Studies Hybridoma Bank for antibodies, and the Bloomington Stock Center for fly stocks. We thank I. Correa for helpful discussions and L. Luo, M. Landgraf, A. Ostrovsky, A. Jenett and members of the Jefferis group for comments on the manuscript. This study made use of the Computational Morphometry Toolkit, supported by the National Institute of Biomedical Imaging and Bioengineering. This work was supported by a European Research Council Starting Investigator Grant, the Medical Research Council (MRC file reference MC_U105188491) and the EMBO Young Investigator Program (GSXEJ); an MRC LMB Graduate Scholarship (JK); a Boehringer Ingelheim Fonds predoctoral fellowship (MJD); a Royal Society Dorothy Hodgkin Fellowship (SC) and an EMBO post-doctoral fellowship (SF).

## AUTHOR CONTRIBUTIONS

JK and GSXEJ initiated the project. JK, JN, SC and GSXEJ designed the experiments. JK, JN, SC and DK made constructs and/or transgenic flies. JK, JN, SC, MJD, BS and SF carried out experiments. JK and GSXEJ analyzed data. JK, SC and GSXEJ prepared figures. JK, JN and GSXEJ wrote the manuscript with input from all authors.

## SUPPLEMENT

**Table S1:** Transgenic fly lines generated in this study. Transgenic fly reporter lines generated in this study are listed. P element insertions were mapped to chromosomes 2 or 3, respectively; insertions into attP sites are indicated in parentheses.

| line | tag | available insertions | description | tag location |
| --- | --- | --- | --- | --- |
| UAS-CD4-CLIP | CLIPf | 2, 3 | plasma membrane targeting | intra |
| UAS-myr-SNAP | SNAPf | 2 (attP40), 3 (attP2) | plasma membrane targeting (myristoylation signal) | intra |
| UAS-myr-TMP | TMP | 2 (attP40), 3 (attP2) | plasma membrane targeting (myristoylation signal) | intra |
| UAS-myr-Halo | Halo | 2 (attP40), 3 (attP2) | plasma membrane targeting (myristoylation signal) | intra |
| UAS-Syt-SNAP | SNAPm | 2, 3 | presynaptic label, fusion to *Drosophila* synaptotagmin (see ref. [ 29]) | intra |
| UAS-Syt-CLIP | CLIPm | 2, 3 | presynaptic label, fusion to *Drosophila* synaptotagmin (see ref. [ 29]) | intra |
| UAS-SNAP-syb | SNAPf | 2, 3 | presynaptic label, fusion to *Drosophila* n-synaptobrevin (see ref. [ 29]) | intra |
| UAS-CLIP-syb | CLIPf | 2, 3 | presynaptic label, fusion to *Drosophila* n-synaptobrevin (see ref. [ 29]) | intra |
| UAS-TLN-SNAP | SNAPm | 2, 3 | postsynaptic label, fusion to mouse ICAM-5 (see ref. [ 31]) | intra |
| UAS-TLN-CLIP | CLIPm | 2, 3 | postsynaptic label, fusion to mouse ICAM-5 (see ref. [ 31]) | intra |
| brp-SNAP | SNAPf | 2 | targeted (MiMIC-based) insertion into *brp* locus | intra |
| brp-SNAP; myrHalo | SNAPf; Halo | 2; 3 (attP2) | targeted (MiMIC-based) insertion into *brp* locus; membrane label | intra |

**Table S2:**
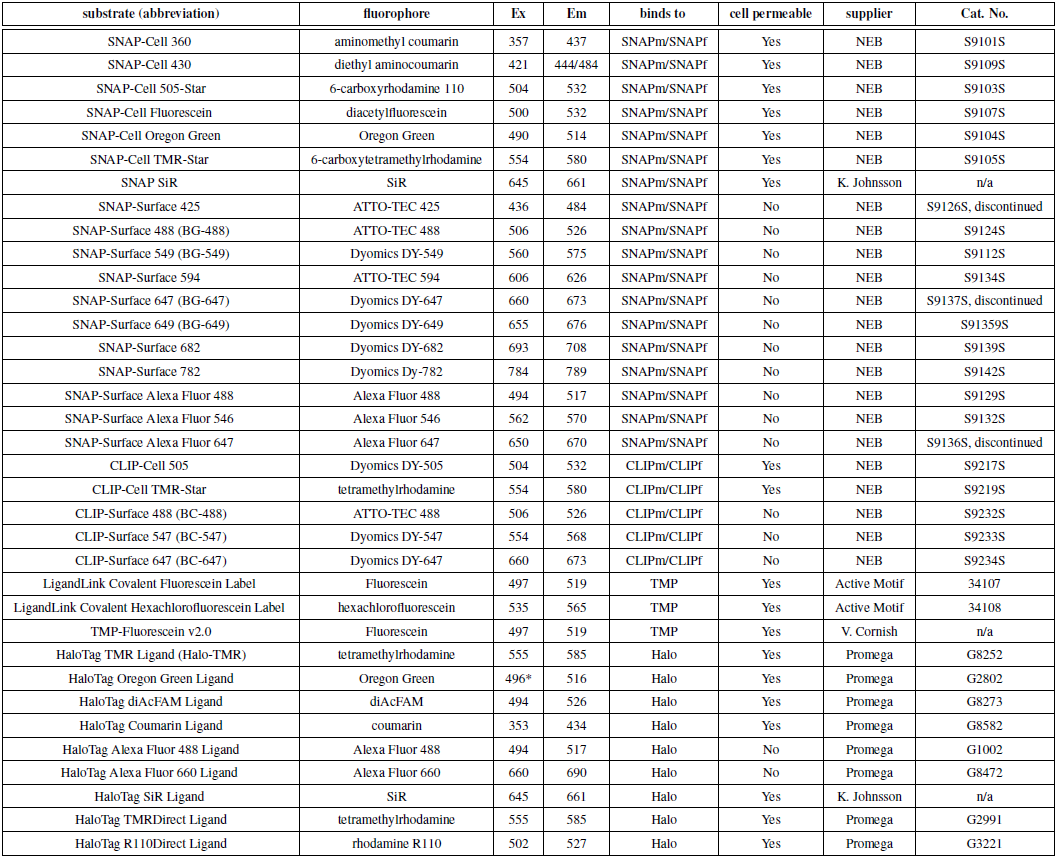
Commercial and custom fluorophore-coupled substrates. Commercially available, fluorophore-coupled substrates for SNAP-, CLIP-, Halo- and TMP-tag are listed.

**Figure S1:**
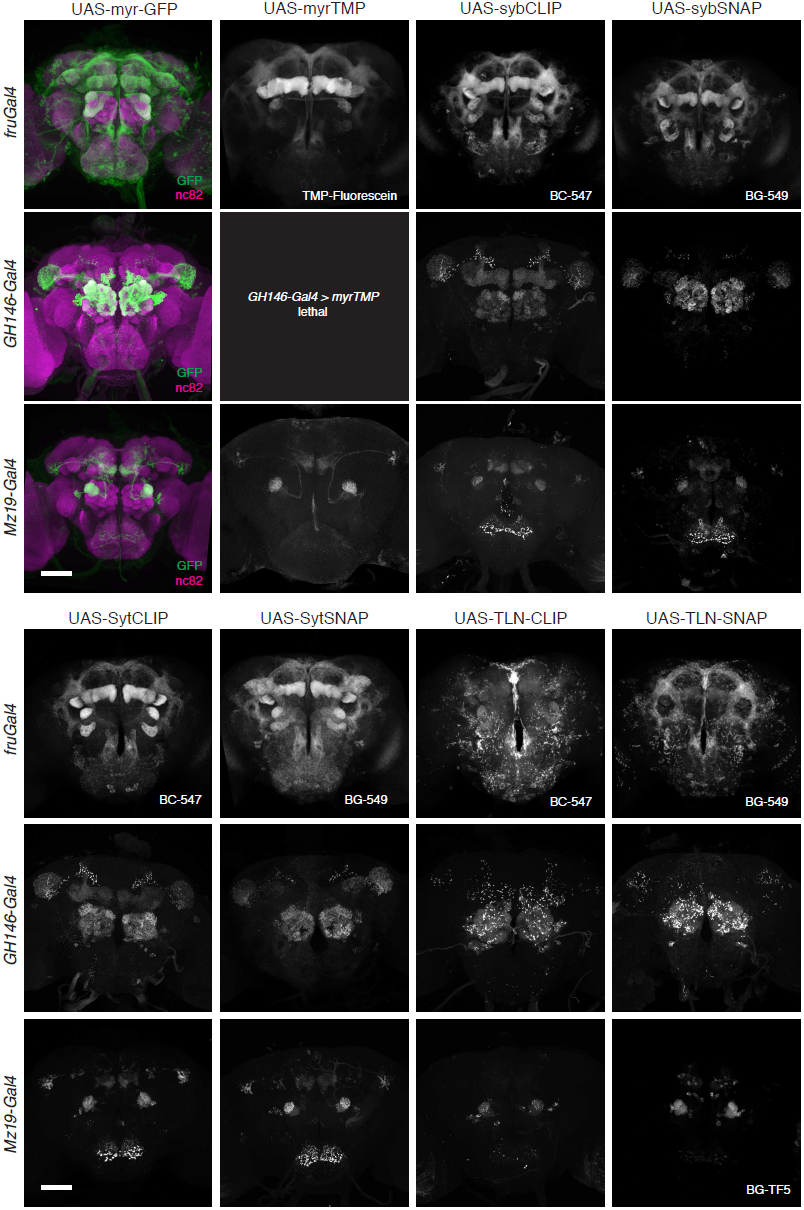
Additional chemical tag reporter fly lines. Expression patterns of additional selected chemical tagging constructs in flies. As in Fig. 1, three different driver lines were used. Images are standard deviation projections from 3D confocal stacks, used substrates are indicated in each image. Scale bars, 50 μm.

**Figure S2:**
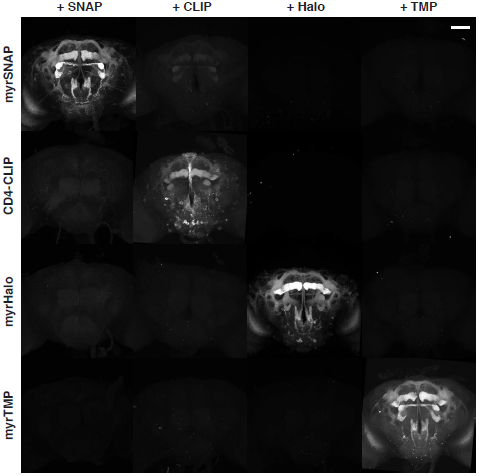
Cross-reactivity of tag-substrate combinations used in this study. Tag-substrate cross-reactivity matrix. Brains in which *fru^Gal4^* drives expression of membrane-bound tags were incubated with matching and non-matching substrates (SNAP, BG-549 1 μM, CLIP, BC-488 1 μM, Halo, Halo-TMR 500 nM, TMP, TMP-Fluorescein 500 nM). No cross-reactivity can be detected, with the exception of CLIP substrate with SNAP-tag. Scale bar, 50 μm.

**Figure S3:**
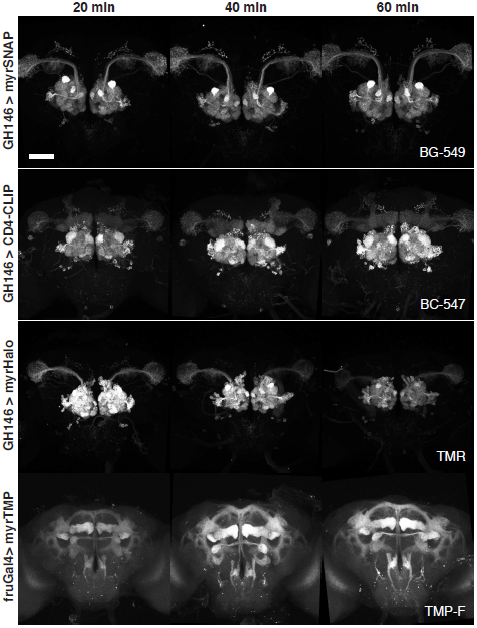
Fixation-resistance of enzyme tags. Brains in which *GH146-Gal4* or *fru*^Gal4^(*GH146-Gal4 > myrTMP* not viable) drive expression of membrane-bound tags were fixed (4% PFA, room temperature) for the indicated time, permeabilized and stained (all substrates 1 μM, 30 min incubation). Note that while SNAP- and CLIP-tag labeling intensities remain largely constant wtih increasing fixation times, Halo-tag labeling intensity decreases and TMP-tag labeling intensity increases. Scale bar, 50 μm.

**Figure S4:**
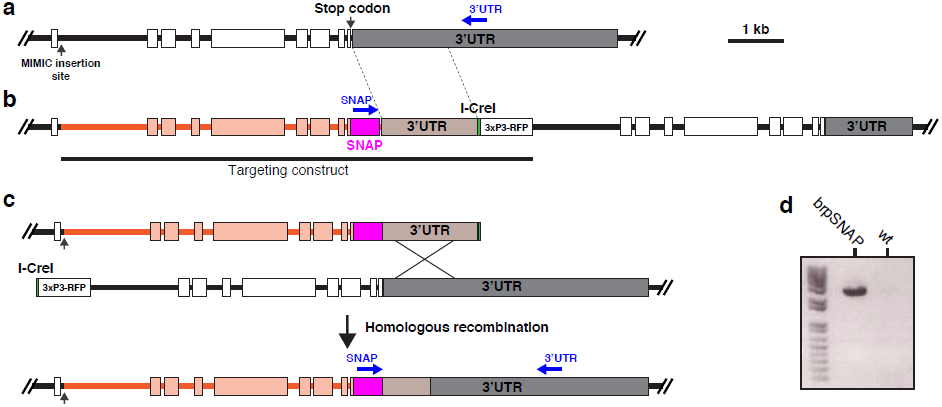
Generation of *brp-SNAP*. Generation of the *brp-SNAP* knock-in allele. (**a**) Schematic showing the organization of the 3’ end of the *brp* locus and the location of the MiMIC insertion. (**b**) Schematic showing the integrated targeting construct and the resulting duplication. (**c**) Resolution of the duplication. The double strand breakage produced by I-CreI is repaired by homologous recombination, resulting in the loss of the RFP marker and the reduction of the duplication. (**d**) PCR with primers ‘SNAP’ and ‘3’UTR’ produces a band in recombinant (brpSNAP), but not wild-type (wt) flies.

**Figure S5:**
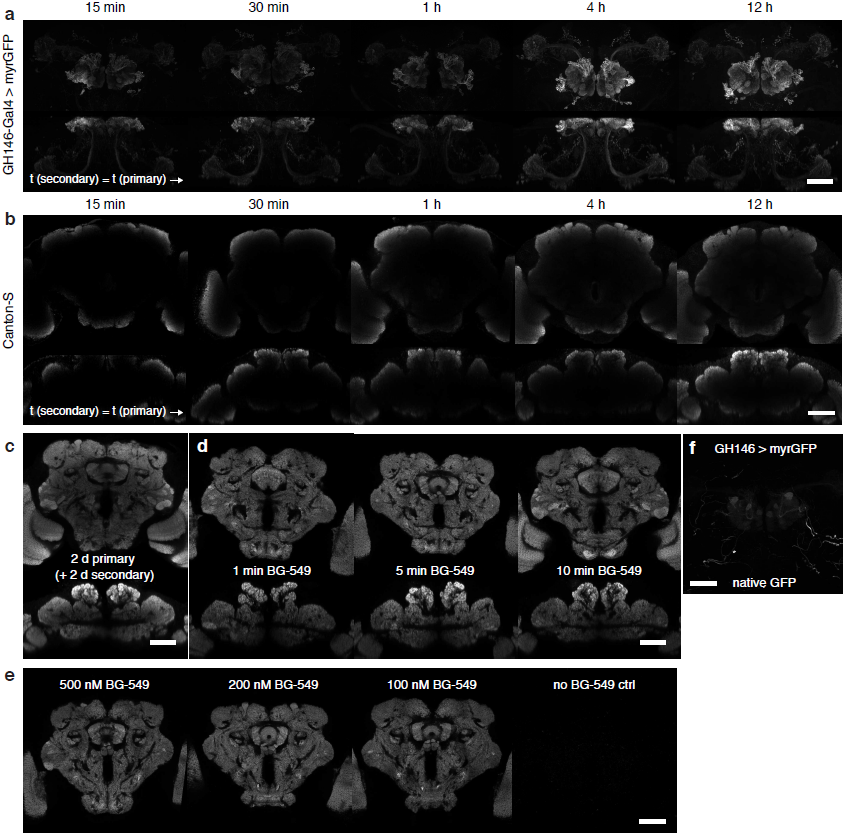
Comparison of neuropil antibody staining and labeling using the *brp-SNAP* transgene. **(a–b)** Staining time course of *GH146-Gal4 > myrGFP* brains with anti-GFP antibody **(a)** and staining time course of the nc82/Bruchpilot neuropil marker using immunostaining against Brp protein **(b)**. (*z-* and *x*-) Maximum projections from 3D confocal stacks are shown in (a), while single coronal (top) and horizontal (bottom) confocal slices through the center of the brain are shown in (b), incubation times are indicated. Note that in contrast to the stainings shown in Fig. 2 (secondary antibodies incubated for 2 d in each case), incubation times are identical for primary and secondary antibodies in (a) and (b). **(c)** Example of nc82-stained brain after full (2 d primary antibody + 2 d secondary antibody) immunostaining protocol. A coronal (top) and a horizontal slice (bottom) from a 3D confocal stack are shown. **(d)** *brp-SNAP* brains after short incubations with BG-549. Coronal (top) and horizontal (bottom) slices through the middle of the brain are shown. Note that homogeneous staining is visible already after 1 min. **(e)** *brp-SNAP* brain stained with lower concentrations of BG-549 substrate and negative control without substrate. Incubation time was 15 min. Coronal slices through the middle of the brain are shown. **(f)** Unstained *GH146-Gal4 > myrGFP* brain imaged after fixation. Most endogenous GFP fluorescence is quenched. Scale bars, 50 µm.

**Figure S6:**
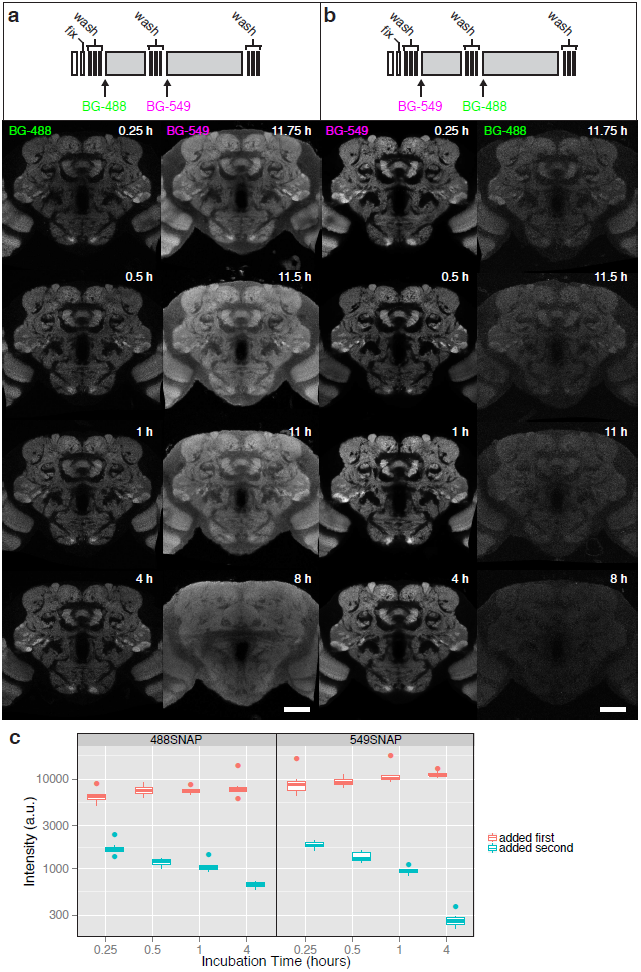
Sequential labeling reveals binding kinetics. Sequential incubation of *brp-SNAP* brains with BG-488 followed by BG-549 (**a**) or BG-549 followed by BG-488 (**b**). The first substrate was added for the indicated time (0.25 h, 0.5 h, 1 h, 4 h), after a washing step the second substrate was added for a total staining time of 12 h. (**c**) Quantification of labeling achieved with BG-488 (left panel) or BG-549 (right panel) using a neuropil mask (see Fig. S7 and methods). Scale bars, 50 μm.

**Figure S7:**
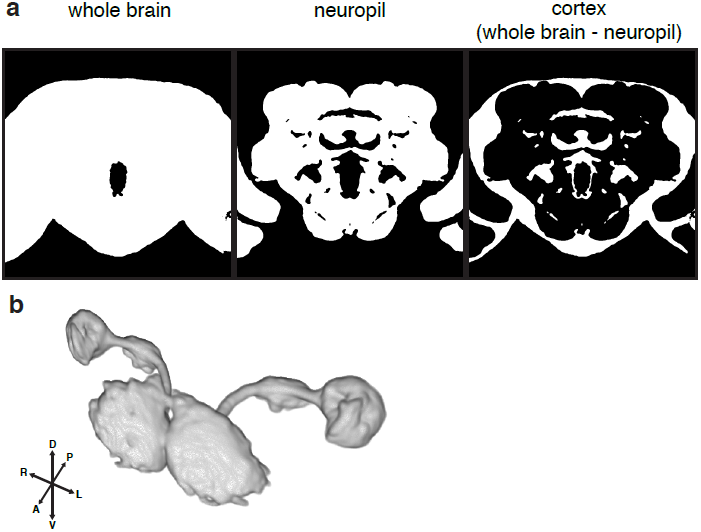
Masks used for quantification of labeling intensity in fly brains. (**a**) Masks used for quantification of labeling in brains stained against nc82- or *brp-SNAP*. Single coronal slices through the 3D masks are shown. (**b**) Volume-rendering of mask used for quantification of labeling in *GH146-Gal4* brains.

